# Visualization of mRNA Expression in *Pseudomonas aeruginosa* Aggregates Reveals Spatial Patterns of Fermentative and Denitrifying Metabolism

**DOI:** 10.1101/2022.03.11.484052

**Authors:** Jadzia Livingston, Melanie A. Spero, Zachery R. Lonergan, Dianne K. Newman

## Abstract

Gaining insight into the behavior of bacteria at the single cell level is important given that heterogeneous microenvironments strongly influence microbial physiology. The hybridization chain reaction (HCR) is a technique that provides *in situ* molecular signal amplification, enabling simultaneous mapping of multiple target RNAs at small spatial scales. To refine this method for biofilm applications, we designed and validated new probes to visualize expression of key catabolic genes in *Pseudomonas aeruginosa* aggregates. In addition to using existing probes for the dissimilatory nitrate reductase (*narG*), we developed probes for a terminal oxidase (*ccoN1*), nitrite reductase (*nirS*), nitrous oxide reductase (*nosZ*), and acetate kinase (*ackA*). These probes can be used to determine gene expression levels both in liquid culture and in biofilms. Using these probes, we quantified gene expression across oxygen gradients in aggregate populations grown using the <u>a</u>gar <u>b</u>lock <u>b</u>iofilm <u>a</u>ssay (ABBA). We observed distinct patterns of catabolic gene expression, with upregulation occurring in particular ABBA regions both within individual aggregates and over the aggregate population. Aerobic respiration (*ccoN1*) showed peak expression under oxic conditions, whereas fermentation (*ackA*) showed peak expression in the anoxic cores of high metabolic activity aggregates near the air-agar interface. Denitrification genes *narG, nirS*, and *nosZ* showed peak expression in hypoxic and anoxic regions, although *nirS* expression was much stronger in anoxic environments compared to other denitrification genes. These results reveal that the microenvironment correlates with catabolic gene expression in aggregates, and demonstrate the utility of HCR in unveiling cellular activities at the microscale in heterogeneous populations.

**Importance:** To understand bacteria in diverse contexts we must understand the variations in behaviors and metabolisms they express spatiotemporally. Populations of bacteria are known to be heterogeneous, but the ways this variation manifests can be challenging to characterize due to technical limitations. By focusing on energy conservation, we demonstrate that HCR v3.0 can visualize nuances in gene expression, allowing us to understand how metabolism in *Pseudomonas aeruginosa* biofilms responds to microenvironmental variation at high spatial resolution. We validated probes for four catabolic genes: a constitutively expressed oxidase, acetate kinase, nitrite reductase, and nitrous oxide reductase. We showed that the genes for different modes of metabolism are expressed in overlapping but distinct subpopulations according to oxygen concentrations in a predictable fashion. The spatial transcriptomic technique described here has the potential to be used to map microbial activities across diverse environments.

## Introduction

Even in an isogenic population, bacteria can display a range of different phenotypes (Ackerman, 2015; Evans, 2020). This phenotypic variation can form along microenvironmental gradients (Lara, 2006) but has also been observed in well-mixed cultures (Thattai, 2004; Kopf, 2015). Heterogeneity in gene expression allows cells to fit different ecological niches and survive in ecosystems with fluctuating conditions, increasing the overall fitness of the population (Thattai, 2004; Smits, 2006; Martins, 2015). Phenotypic heterogeneity is relevant in a variety of contexts, from rhizosphere colonization (Broek, 2005) to metabolic engineering for optimal biosynthesis of fuels and medicines (Xiao, 2016). In chronic infections, phenotypic variation shields pathogens from antibiotics, allowing subpopulations with distinct metabolic activities to survive rounds of treatment (Michiels, 2016; Fisher, 2017) and contribute to the development of antibiotic resistance (Cohen, 2013).

While many technologies exist to study gene expression at the population (Guell, 2011) and single cell (Kuchina, 2020) levels, few allow the mRNA output of individual bacteria to be observed while maintaining the microenvironmental context and without genetically modifying the cells (e.g. use of genetically-encoded fluorescent proteins). Recently, a spatial transcriptomic method was introduced that enables visualization of hundreds of transcripts at a time in single cells (Dar, 2021), yet this method requires specialized instrumentation and analysis software, providing a barrier to widespread usage. By contrast, the hybridization chain reaction (HCR) is a more accessible technology that permits single cell transcript visualization of a smaller number of genes using standard instrumentation (Choi, 2018; Trivedi, 2018). HCR is a form of fluorescence *in situ* hybridization (FISH), where the binding of fluorescent probes to a target RNA of interest results in the binding of additional fluorescent probes, thus amplifying the signal (Choi, 2018). The technology first was developed in eukaryotes (Dirks, 2004), but has been used in bacteria to image rRNA for the purposes of species identification and growth rate assessment in mixed communities (Nikolakakis, 2015; Depas, 2016), as well as to visualize gene expression in heterogeneous populations (Jorth, 2020).

Building on these studies, we reasoned that HCR was well suited to enabling a systematic exploration of the variation in catabolic activities within aggregate biofilms of the model biofilm forming bacterium *Pseudomonas aeruginosa*. One environmental parameter that has profound effects on the metabolic state of bacteria is oxygen. For example, many infection sites have large regions of hypoxia, including those found in skin wounds, lungs, and the gut (Palmer, 2007; Hong 2014), Additionally, opportunistic pathogens like *P. aeruginosa* can form aggregate biofilms during infection, (Grant, 2013; Zhao, 2013; Depas, 2016), and biofilms produce their own oxygen gradients, with biofilm-interior populations experiencing oxygen limitation (Stewart, 2003). This has devastating consequences for human health because commonly used antibiotics such as tobramycin and ciprofloxacin are ineffective at killing oxygen-limited *P. aeruginosa* at the center of biofilms, while drugs that target anaerobic metabolism specifically kill oxygen-limited populations (Walters, 2003; Spero, 2018).

Accordingly, antibiotic treatments must take the microenvironment and the phenotypic heterogeneity experienced by opportunistic pathogens in the host into account to be successful (Spiro, 2012; Cook, 2015). Before we can rationally design better treatments, we must improve our understanding of the range of metabolic states that exist in biofilm populations spatiotemporally (O’Toole, 2021).

In the absence of oxygen, *P. aeruginosa* cells utilize different strategies to conserve energy, ranging from oxidative to substrate-level phosphorylation (Schobert, 2010). Because chronic infections are often oxidant limited, how *P. aeruginosa* responds to electron acceptor availability is expected to lead to metabolic differentiation within these environments, which partly underpins its antibiotic tolerance and resistance (Ciofu, 2019). Under anoxic conditions, *P. aeruginosa* can use nitrate and other N-oxides as alternative terminal electron acceptors to oxygen by performing denitrification (Arat, 2015). The nitrate levels in chronic surgical wound and cystic fibrosis lung infections are high enough to support denitrification in some regions, while the oxygen levels are low enough to induce denitrification (Palmer, 2007; Line, 2014; Turner, 2014). In the absence of nitrate, *P. aeruginosa* may survive under anoxic conditions for long periods by performing pyruvate fermentation, in which pyruvate is converted to lactate and acetate (Eschbach, 2004), or by fermenting arginine (Vander Wauven, 1984) or using phenazines to facilitate glucose fermentation (Glasser, 2014). Aerobic, denitrifying, and fermentative metabolisms are all relevant to bacterial populations in chronic infection environments.

If we are to use HCR to map these metabolisms within biofilm aggregates, it is important to interpret any gene expression patterns we might see within an ecophysiological regulatory framework. In *P. aeruginosa*, the primary respiratory oxidase under highly aerobic conditions is cbb_3_-1, encoded by *cco1* (Jo, 2014). *cco1* is constitutively expressed when oxygen is replete but slightly downregulated in stationary phase (Alvarez-Ortega, 2007; Arai, 2011). Denitrification is catalyzed by a series of reductases, in particular *nar, nir, nor*, and *nos* (Arai, 2011; Arat, 2015).

The main regulatory proteins that govern the expression of these genes are Anr, Dnr, NarXL, and NirQ (Figure 1). Under anoxic conditions, Anr activates Dnr, a regulatory protein that is downregulated by phosphate availability, which activates NirQ. Where nitrate is available and oxygen is limited, Anr additionally activates NarXL which in turn amplifies the activation of Dnr and NirQ. The nitrate reductase gene, *nar*, is upregulated by NarXL, and this upregulation is amplified by Dnr in the presence of nitric oxide (Arat, 2015). The nitrite reductase gene, *nir*, is highly upregulated under anoxic conditions and has higher differential expression from oxic to anoxic conditions than *nar* (Palmer, 2007; Alvarez-Ortega, 2007; Arai, 2011; Arat, 2015). *nir* is slightly upregulated by Anr, causing a small amount of nitrite to be converted to nitric oxide, which triggers Dnr and increases the upregulation of *nir* in a feedback loop (Kuroki, 2014). NirQ and Dnr tightly coregulate *nir* with the nitric oxide reductase *nor* in response to nitric oxide availability, preventing nitric oxide from accumulating (Arat, 2015). The nitrous oxide reductase, *nos*, is upregulated by Dnr in response to nitric oxide availability (Arat, 2015). In contrast, the acetate kinase gene, *ackA*, responsible for ATP synthesis during pyruvate and phenazine-mediated glucose fermentation, is induced by the regulatory proteins Anr and IHF (Integration host factor).

**Figure 1:**
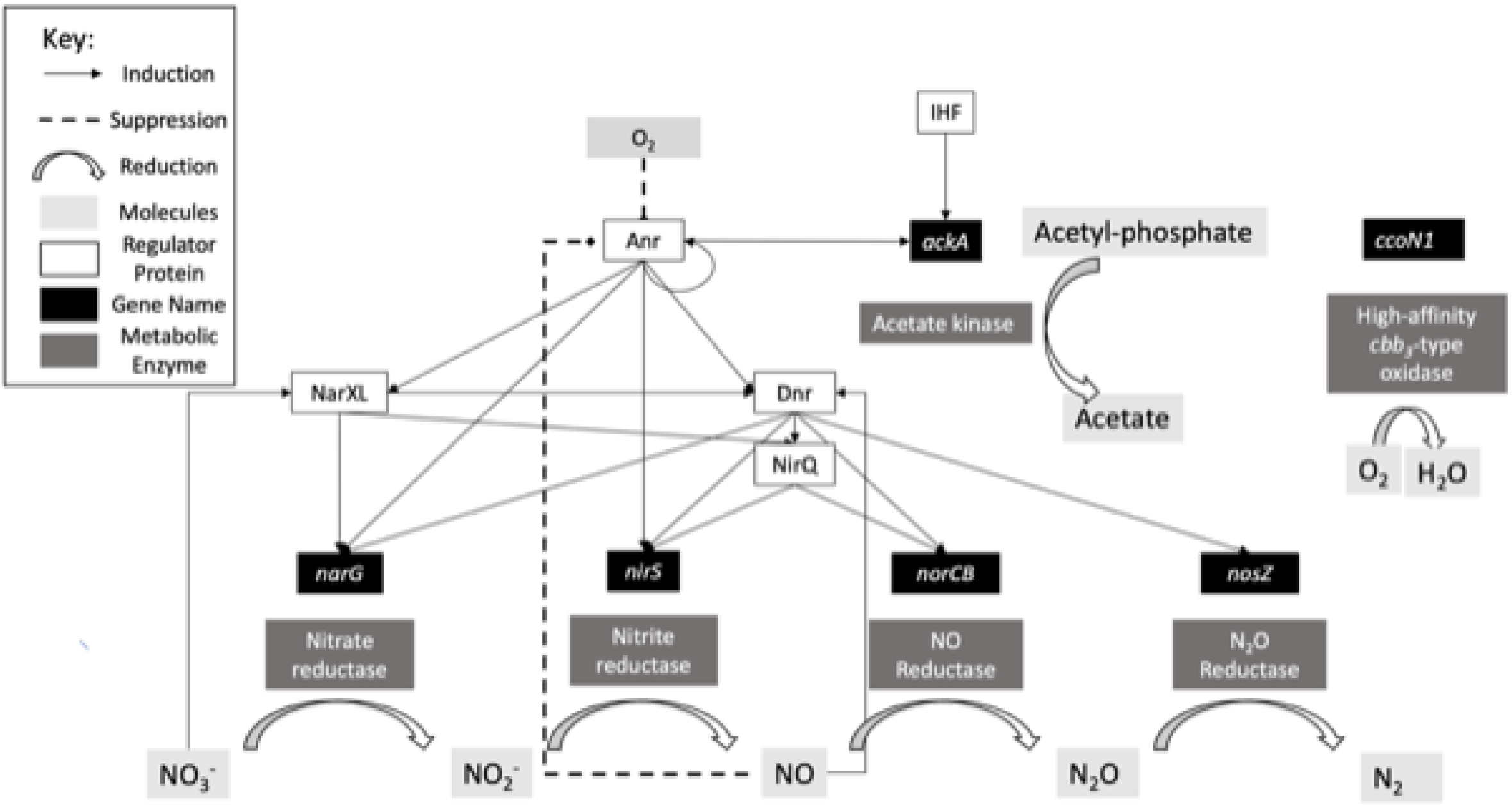
Regulatory Diagram of Target Genes. This diagram was compiled from a literature search of transcriptomics reviews in *P. aeruginosa*. (Eschbach, 2004; Arai, 2005; Schreiber, 2007; Arai, 2011; Arat, 2015)

We targeted the expression levels of *cco1, narG, nirS, nosZ* and *ackA* mRNA molecules to investigate the spatiometabolic patterning of catabolic gene expression within and across *P. aeruginosa* aggregate biofilm populations. Our goal was to use a finite set of genes to identify the range of metabolic states adopted by multicellular aggregates under oxidant-limited conditions. Here we show that HCR 3.0 not only enables spatiometabolic mapping *in situ*, but that it allows us to infer the metabolic state and environmental conditions that single cells within these heterogeneous populations are experiencing.

## Results

### HCR Probes are Specific for their mRNA Targets

To validate the probes designed for this study, we tested their specificity using WT and mutant strains in which the target gene had been cleanly deleted. To test *narG* and *nirS* probes, we grew WT, Δ*narGHJI*, and Δ*nirS* strains aerobically in LB supplemented with 40 mM nitrate to late exponential phase to achieve oxygen limitation and activate Anr. To test *ccoN1* probes, we grew WT and Δ*ccoN1* strains aerobically in the same medium to early exponential phase to achieve highly oxygenated conditions. To test *nosZ* probes, we grew WT and Δ*nosZ* strains anaerobically in the same medium to mid-exponential phase to achieve denitrifying metabolism. We were unable to observe *ackA* signal in any of the liquid culture conditions we tested, so to test *ackA* probes, WT and Δ*ackA* were grown in the agar biofilm block assay (ABBA) for 12 hours in LB medium and 0.5% agar, supplemented with 40 mM nitrate. Briefly, in ABBA, *P. aeruginosa* cells are suspended in an agar medium that is allowed to solidify. Cells then grow as aggregate biofilms suspended in this agar matrix, and nutrient gradients are known to develop within individual aggregates as well as across the population of aggregates (see below).

The *narG, nirS, nosZ, and ccoN1* probes showed an average mean intensity per cell that was an order of magnitude higher in the wildtype compared to the deletion strains, indicating our probe sets are specific for each target mRNA (Figures 2A, 2C). Additionally, the *ackA* probe showed significantly higher average mean intensity per ABBA aggregate compared to the deletion strain (Figures 2B, 2D). In some of the deletion mutant controls, particularly for the *nosZ* and *ccoN1* probe sets, we occasionally observed fluorescent puncta that did not overlap with cells and appeared qualitatively different from the signal observed in the positive control. We therefore assume that these puncta were due to non-specific binding to the slide and not biologically significant.

**Figure 2:**
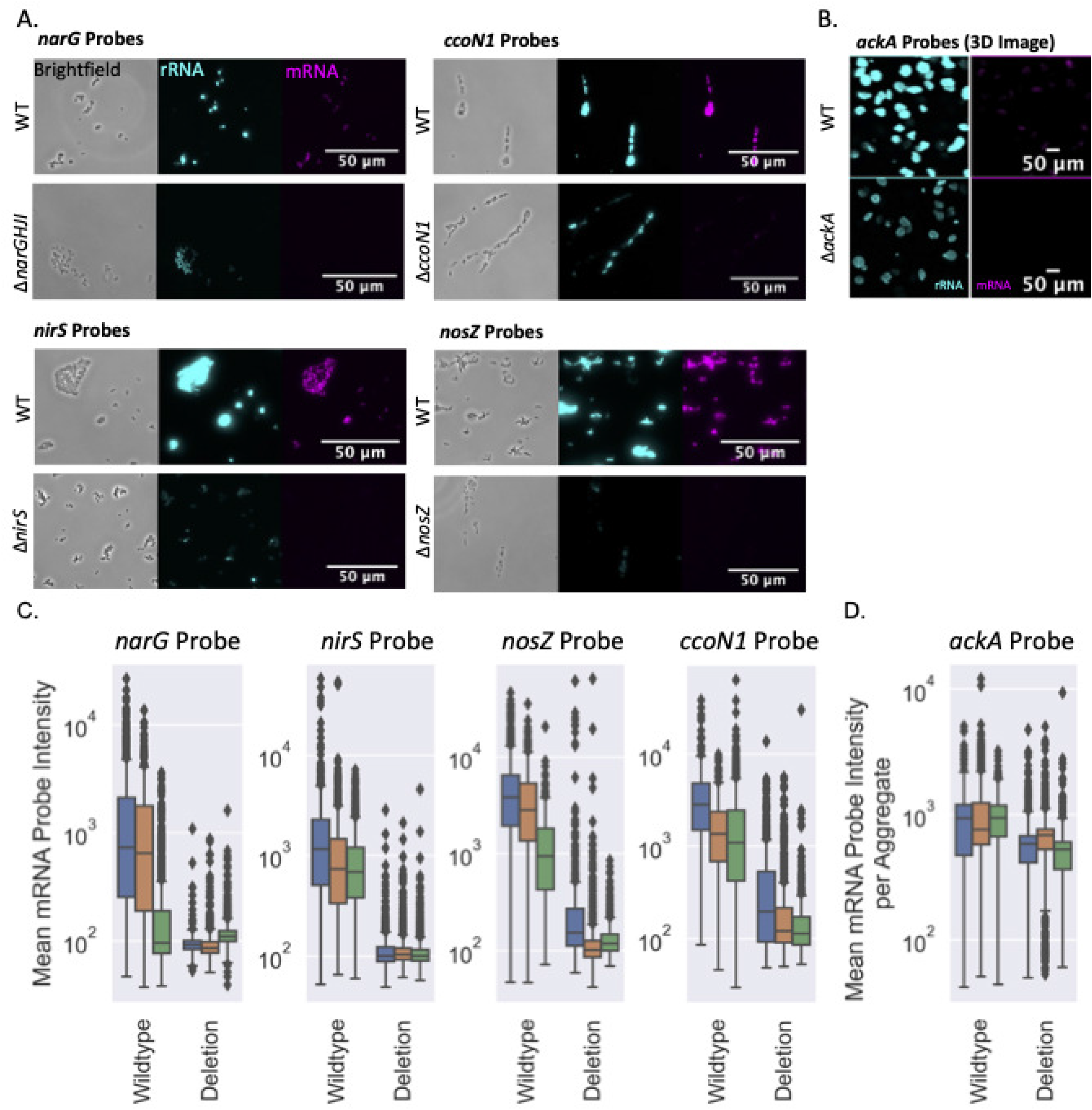
mRNA Probes Are Target-Specific. (A) Micrographs of Single Cell Deletion Validation Controls for mRNA Probes. (B) Micrographs of Aggregate Deletion Validation Controls for mRNA probes. Images reflect the region 50-100 μm from the air-agar interface. (C) Quantification of mean mRNA probe intensities for single cells grown in liquid culture of wildtype and deletion mutants for target genes under upregulating growth conditions. Mean intensity was 10-fold higher in the wildtype than in the deletion for all probe sets. Each boxplot summarizes approximately 10 images per replicate. Three replicates were performed per condition. (D) Quantification of mean mRNA probe intensities for aggregates grown in agar blocks wildtype and deletion mutants for *ackA* under upregulating growth conditions. Mean intensity was 2-fold higher in the wildtype, and each boxplot summarizes 3-5 images per replicate. Three replicates were performed per condition.

### Metabolite Gradients are Readily Established in ABBA

To explore the phenotypic heterogeneity of *P. aeruginosa* wildtype PA14, we used HCR to visualize the localization of *narG, nirS, nosZ, ccoN1*, and *ackA* expression across biofilm aggregates grown in the ABBA. Within ABBAs, cells generate oxygen and nitrate gradients over the course of a few hours. We used a thresholded 16S rRNA intensity value to define aggregate boundaries and to control for baseline metabolic activity, and quantified mRNA and rRNA signals within this mask. *P. aeruginosa* aggregates were grown in LB medium with 0.5% agar, supplemented with 40 mM nitrate. We chose these conditions to capture the effects of oxygen and nitrate gradients on the expression of our genes of interest. Physiological levels of nitrate in infected cystic fibrosis lung samples and chronic wounds are ∼400 μM (Palmer, 2007; Bernatchez, 2013), but we chose a higher concentration of nitrate to ensure that it would still be available after 12 hours of growth.

In an ABBA, an oxygen gradient develops, and the oxygen minimum migrates upwards towards the air-agar interface over the course of the incubation as cell density increases, and thus oxygen availability decreases (Figure 3A). The exact depths at which oxygen is minimal vary between agar blocks, but the pattern is conserved. A steep oxygen gradient occurs across the first ∼100 microns directly below the surface, and regions deeper than ∼250 μm are uniformly oxygen starved. Accordingly, deeper aggregates in the anoxic zone will have experienced anoxia for longer than the aggregates above them. As aggregates begin experiencing hypoxia, cells will upregulate denitrification genes and reduce nitrate, such that a similar nitrate gradient is also expected to develop. The deepest aggregates are likely the first to begin consuming nitrate in their local environment, so deeper aggregates likely have less access to nitrate at this time point. As the denitrification pathway proceeds, nitric oxide is generated (Williams, 1978), so a greater amount of nitric oxide is expected at lower depths.

**Figure 3:**
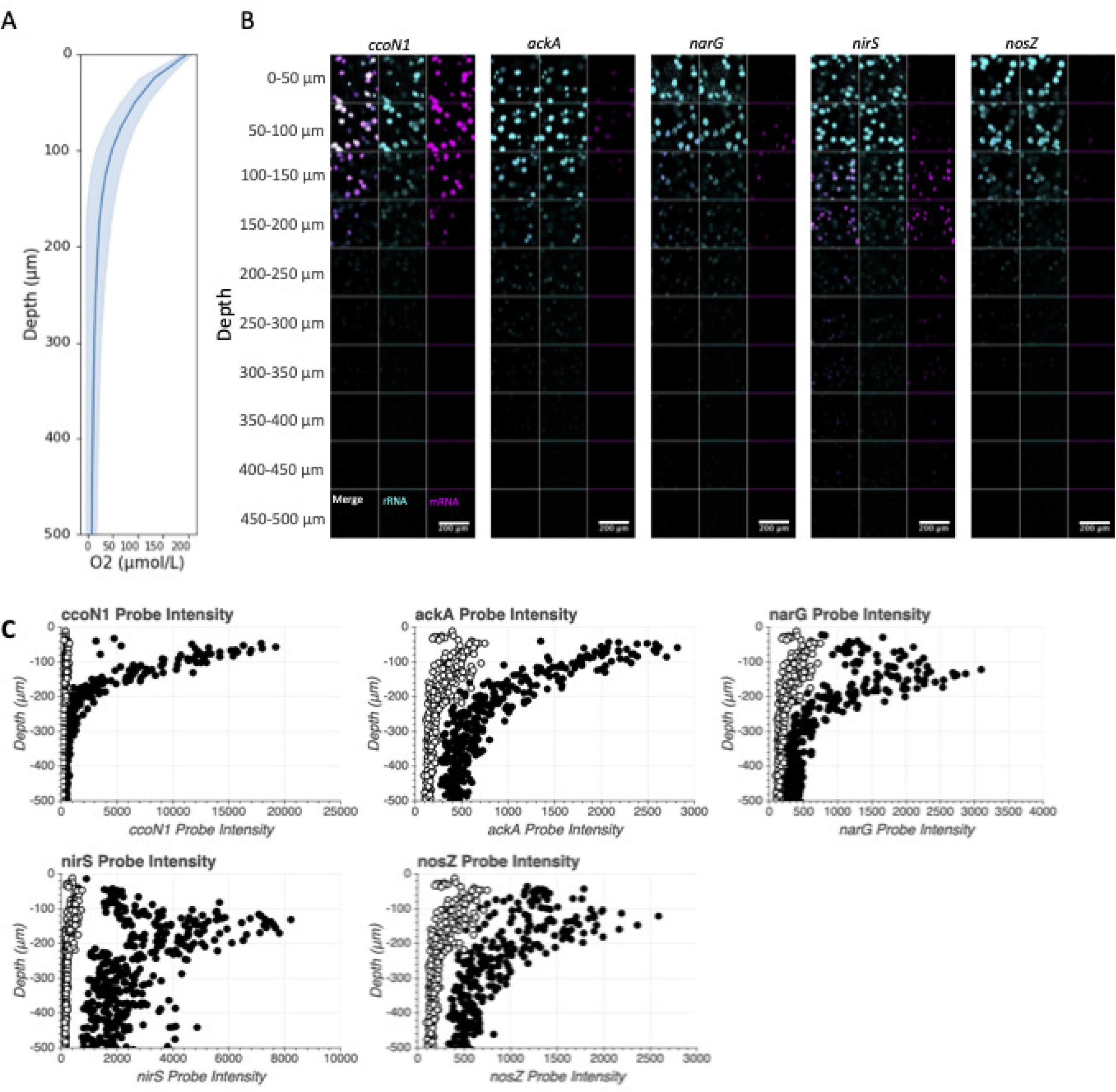
Metabolic Genes Show Distinct Patterns Across 3D Oxygen Gradients. (A) Mean oxygen levels of ABBA samples grown with LB+40 mM Nitrate for 12 hours. The dark line is the mean, while the shading represents the standard deviation of 8 biological replicates grown under identical conditions. (B) 3D Micrographs of Probe Signal in LB + 40 mM Nitrate ABBAs. Each image represents a 50 micron slice of agar, compiled from 8 individual z-slices with an interslice distance of 6.24 microns, viewed from the top of the block, with each sequential image from the top of the figure representing the section directly below the slice above it. rRNA signal is colored cyan, while mRNA signal is colored magenta. (C) Mean mRNA Channel Intensity per aggregate (x axis) plotted by depth. Each plot represents four images of one replicate each of an experimental and control condition, and each point represents one aggregate. Filled points represent the experimental condition, while the white circles represent a control condition where only rRNA probes were used.

Given these conditions and the regulatory architecture controlling expression of these genes (Figure 1), we predicted that *narG* would be expressed in regions with high nitrate and low oxygen concentrations, *nirS* would be expressed in regions with high nitrate and/or nitric oxide and low oxygen concentrations, and *nosZ* would be expressed in regions with nitric oxide present and low oxygen concentrations. We expected *ccoN1* to be expressed most highly in regions with high oxygen and *ackA* to be expressed in regions that are limited for both oxygen and nitrate.

### Metabolic Genes Show Distinct Expression Patterns within ABBAs

The oxidase *ccoN1* was maximally expressed in the oxic region (0-50 μm from the air-agar interface at the top of the ABBA blocks), and steeply decreased in expression deeper into the block along the oxygen gradient which is the same pattern shown by the 16S rRNA signal (Figure 3B). In the larger aggregates closer to the surface, signal for *ccoN1* and 16S rRNA was higher along the aggregate exterior than in the centers, while signal for the *ackA* gene was highly expressed in the interior (Figures 3B,3C, Figure S1). Elevated signal for the denitrification genes was not detected in this region.

In the region below the oxic portion of our experimental system (>50 μm from the air-agar interface at the top of the ABBA blocks), rRNA, *ccoN1* and *ackA* expression was reduced, and expression of the denitrification genes were elevated, correlating with low oxygen concentrations that decreased with depth across this region (Figure 3A, 3B, 3C) All three denitrification genes were expressed at their maximal levels in aggregates in this region. The nitrate reductase *narG* was upregulated under hypoxic conditions in the 40 mM nitrate samples, peaking in average mean intensity per aggregate at a depth of 100-150 μm from the air-agar interface (Figure 3B, 3C). *narG* signal was restricted to the interior of aggregates, where oxygen availability is expected to be reduced compared to cells on the aggregate exterior (Figure 3B, Figure S1). The nitrite reductase *nirS* was upregulated under hypoxic and anoxic conditions, peaking between 100-200 μm from the surface. The nitrous oxide reductase *nosZ* was expressed in cells in the cores of some aggregates in the hypoxic and anoxic regions, but was not uniformly expressed by aggregates at any depth (Figure 3C). Oxygen levels reached their minimal levels between 150-250 μm from the air-agar interface (Figure 3A).

Oxygen levels were uniformly low below 250 μm from the air-agar interface. All genes measured showed reduced expression with depth across this anoxic region, likely due to oxidant limitation, although *nirS* expression was elevated in the cores of aggregates at all depths. *nirS* signal was restricted to the interior of aggregates in the hypoxic zone, but was expressed across entire aggregates further into the anoxic zone (Figure 3B, Figure S1).

Appreciable *narG* signal was not detected in the deepest anoxic region (Figure 3C). Data from two additional replicates of this experiment (mRNA and rRNA ABBA images with quantification) can be found in Figures S2-S7.

### Differentially Regulated Metabolic Genes show Different Expression Patterns than rRNA

By comparing the rRNA and mRNA signal per aggregate in each sample, we can detect where the mRNA signal diverges from the basal metabolic rate, reflected by the rRNA signal. The *ccoN1* signal linearly and positively covaried with the rRNA signal across depths (Figure 4). No other genes showed this pattern. All three denitrification genes showed maximal expression in regions where the rRNA signal was ≤ half the maximum, corresponding to the hypoxic transition region of the agar block. The *nirS* signal showed the highest negative covariance with the rRNA signal, with maximal expression occurring in aggregates with one-third the maximum rRNA signal. The *ackA* signal showed positive, but non-linear, covariance with the rRNA signal, showing the highest mRNA signal in the aggregates at the surface with the highest rRNA signal, and linearly covarying throughout the oxic and hypoxic regions before a steep dropoff of mRNA signal in the anoxic region. The rRNA and *ccoN1* signals were qualitatively highest at the aggregate edges, while the *narG, nirS*, and *nosZ* signals were highest in the aggregate cores (Figure S1), likely indicating a radial oxygen gradient from the edges to the cores of individual aggregates. The *ackA* signal was heterogeneously expressed in some cells throughout aggregates.

**Figure 4:**
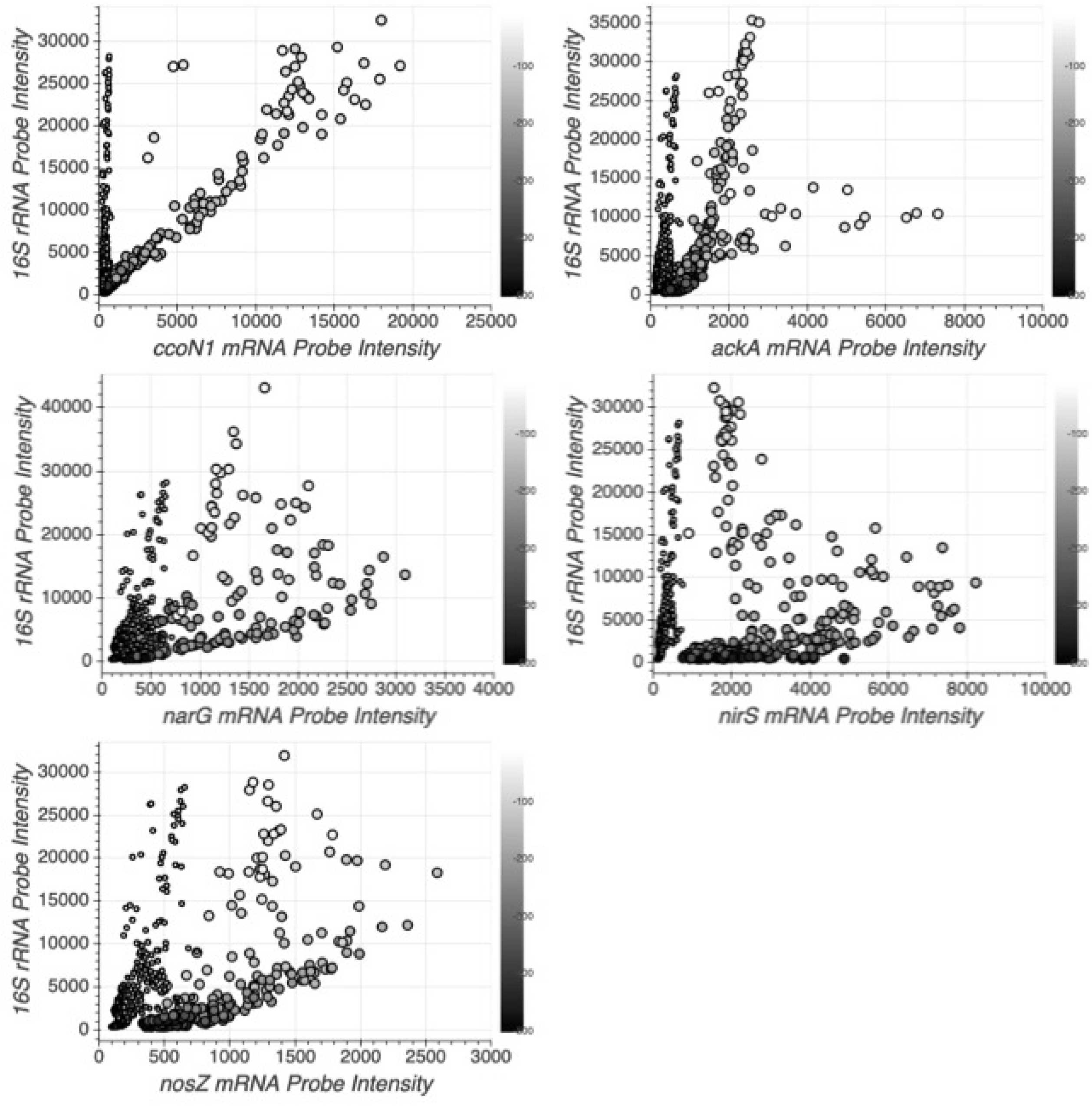
Metabolic Genes Show Distinct Patterns Compared to rRNA Expression. Mean mRNA intensity per aggregate (x) plotted against mean rRNA intensity per aggregate (y) colored by depth. Each plot represents four z-stacks of one replicate, and each point represents one aggregate. Large points represent the experimental condition, and small points represent the control condition without mRNA probes.

## Discussion

In this work, we developed and validated a set of HCR probes targeting *mRNA* molecules for key denitrification, aerobic respiration, and fermentation genes in *P. aeruginosa*. This targeted probe set permits mapping of the metabolic states of cells in aggregate biofilms at high resolution across microenvironments characterized by steep oxygen and nitrate gradients. Our results reveal distinct patterns of catabolic gene expression within these populations, underscoring that metabolic activity can be heterogeneous over small spatial scales. The demonstration that HCR can be applied to chart catabolism within a reductionist laboratory biofilm sets the stage for future studies of microbial responses to dynamic microenvironments in more complex contexts. Importantly, HCR permits the study of gene expression dynamics in complex wildtype populations without the need for generating reporter strains; though developed here for a model organism, this capability has obvious advantages for the study of strains that are not yet genetically tractable.

Our results show that the relationship between *ccoN1* and rRNA signals for entire aggregates is linear and positive, indicating that changes in *ccoN1* expression reflect changes in the metabolic rate on the population level (DePas, 2016). This is consistent with previous studies showing *ccoN1* as constitutively expressed (Jo, 2017). Most genes in Gram-negative bacteria such as *P. aeruginosa* show basal levels of gene expression even outside the upregulation condition (Volke, 2020) so the expression pattern shown by *ccoN1* represents the null hypothesis of what expression should look like if the gene is not regulated by environmental conditions. If gene expression patterns diverge from the pattern shown by *ccoN1* and rRNA, we can infer that they are upregulated and downregulated based on additional conditions other than the metabolic rate.

We observed that expression of the genes encoding the denitrification machinery in *P. aeruginosa* is spatially stratified across the ABBA system after 12 hours. Previous studies have demonstrated that denitrification in liquid cultures occurs in successive steps over time (Williams, 1978). Recently this sequential pattern was observed in structured communities of *P. aeruginosa* for *narG* and *nirS* using reporter strains (Smirga, 2021), revealing maximal expression of these genes moved outward from the center of particles suspended in bulk medium across time, suggesting aggregates progressively extinguished local oxygen, nitrate, and nitric oxide concentrations. Our findings confirm this spatial pattern in *P. aeruginosa*, showing that at a single time point maximal *narG* expression occurs closer to the air-agar interface than maximal *nirS* and *nosZ* expression, as oxygen limitation and nitrate limitation increase with depth. Furthermore, we were able to visualize this spatial pattern in individual aggregates. In aggregates between 50-150 μm from the air-agar interface, *narG* expression occurs closer to the exterior of individual aggregates compared to *nirS* and *nosZ* expression, and *nirS* expression is present across entire aggregates at greater, mostly anoxic depths but limited to the cores of aggregates in hypoxic conditions.

Despite both being upregulated by Anr, fermentation and denitrification genes showed distinct expression patterns. The fermentation gene *ackA* was upregulated in the cores of the aggregates closest to the air-agar interface where oxygen concentrations are higher, while the denitrification genes were upregulated in the cores of the aggregates in the hypoxic regions.

Both types of metabolism mark hypoxic/anoxic conditions, but they are regulated in response to distinct nitrate and nitric oxide conditions. Given that *ackA* is expressed in the absence of nitrate (Eschbach, 2004), the pattern of the denitrification and fermentation genes suggests that the cores of the aggregates at the top of the agar blocks may be conserving energy by fermentation, due to a microenvironmental regulatory cue. Notably, the denitrification genes are directly upregulated by Dnr, which is activated by nitric oxide, while *ackA* is directly upregulated by Anr, which is deactivated by nitric oxide. We can infer from this regulatory relationship that nitric oxide signaling is occurring in the lower aggregates but not in the upper aggregates at this time point. Direct measurement of N-oxide concentrations in the ABBA system is important for understanding the relationship between microenvironmental conditions and metabolic state, and will be the focus of future experiments.

Going forward, this set of probes may be expanded and multiple high-signal probes may be used to determine correlation and anticorrelation more precisely. HCR may be used in two directions: read-out, in which specific regions of a sample are queried to learn which genes are co-expressed within them, and read-in, in which knowledge of genes whose co-expression is already understood is leveraged to infer something about the microenvironment of the region in which they are expressed (Trivedi, 2018). The former application allows us to choose a specific context in space and time and see which genes are being expressed without prior knowledge, whereas the latter approach allows us to apply our knowledge of gene regulation to infer something about the microenvironment–such as we did in this study to dissect the microenvironmental conditions of ABBA subpopulations. In the future, the probes developed in this study may be applied to observe metabolisms and infer potential microenvironmental conditions experienced by *P. aeruginosa* in clinical samples, opening the doors for greater insight into the metabolic processes sustaining cells in infections.

## Acknowledgements

We thank Lars Dietrich and Lindsey Christine Florek for the Δ*ccoN1* PA14 strain, Steven Wilbert for the Δ*nirS* and Δn*osZ* strains, and David Basta and Nate Glasser for the Δ*ackA* strain. We thank Chelsey VanDrisse, Steven Wilbert, John Ciemnieki and Georgia Squyres for constructive feedback and we thank Niles Pierce, and Maayan Schwarzkopf for assistance using the HCR system. Confocal microscopy was performed in the Caltech Biological Imaging Facility at the Caltech Beckman Institute, which is supported by the Arnold and Mabel Beckman Foundation, and we thank Giada Spigolon for her help using the facility. This work was supported by grants from the NIH (R01 HL152190-03) to DKN, the Doren Family Foundation, the Cystic Fibrosis Foundation (postdoctoral fellowship to MAS, SPERO19F0), and the Jane Coffin Childs Memorial Fund (postdoctoral fellowship to ZRL).

## Materials and Methods

### Microbial strains and growth conditions

*P. aeruginosa* strains UCBPP-PA14 (wild type (WT)) and isogenic Δ*narGHJI* (Spero, 2018), Δ*ccoN1* (Jo, 2017), Δ*ackA* (Glasser, 2014), Δ*nirS* and Δ*nosZ* (Wilbert, 2021) were routinely grown in 5 mL Luria Bertani (LB) broth supplemented with 40 mM KNO3 shaking at 37°C. Aerobic cultures were incubated at 37°Cshaking at 250 rpm unless described otherwise. Anaerobic cultures were incubated in an anaerobic chamber at 37°Cwithout shaking.

### HCR v3.0 probe validation

WT and deletion mutant cultures were grown overnight from freezer stocks in LB broth supplemented with 40 mM KNO3. For the *ackA* mutant, cells were grown in the agarose biofilm block assay (ABBA) as described below. For other mutants, 20 μL overnight culture was inoculated into 5 mL of pre-warmed and degassed media and incubated under conditions meant to induce upregulation of the target gene. For *narG* and *nirS*, cultures were grown aerobically as described above for 4 hours to an approximate optical density at 500 nm of 1.1. For *ccoN1*, cultures were grown aerobically as described above for 2.5 hours to an approximate optical density at 500 nm of 0.2. For *nosZ*, wildtype and deletion mutant cultures were grown anaerobically for 2.5 hours, to an approximate optical density at 500 nm of 0.2. 1 mL of culture was pelleted and cells were fixed by resuspending in 1 mL of 4% paraformaldehyde in phosphate-buffered saline (PBS) pH 7.2, and incubating resuspensions at 25°C for 90 minutes. After fixing, cells were pelleted, washed with PBS, resuspended in 1mL of a 1:1 ethanol:PBS solution, and stored at −20C until HCR 3.0 experiments were performed.

Prior to performing HCR, cell suspensions were washed in PBS and treated with 1 mg/ml lysozyme (catalog number L6876; Sigma) in 10 mM Tris-HCl (pH 7.6) for 1 h at 37°C with shaking. After these treatments, cells were washed with PBS, resuspended in 1mL of a 1:1 ethanol:PBS solution. Cell solutions were suspended in hybridization buffer for 1 hour at 37°C. Probe solutions were prepared, containing 5 nM of appropriate HCR v3.0 initiator odd and even probe pairs in filter-sterilized hybridization buffer. 20 uL of probe solutions (Integrated DNA Technologies) were added to each cell solution. *Pseudomonas aeruginosa* rRNA and *narG* mRNA sequences were targeted using target sequences previously used in Jorth, 2020. Probes specific to *nirS, nosZ, ackA*, and *ccoN1* were designed in the study, and the target sequences are included in Table S1. Solutions were hybridized overnight at 37°C, then washed twice with 200 uL probe wash buffer at 37°C. Cells were then resuspended in 40 μL amplification buffer and pre-amplified for 30 minutes, while fluorescent hairpin probes (Molecular Technologies) were thawed, each denatured in separate tubes in a thermal cycler for 90s at 95°C and allowed to cool for at least 30 m at 25°C in the dark.

Hairpins with Alexa Fluor 488 were used for the rRNA probes, and hairpins with Alexa Fluor 647 were used for the mRNA probes. The B1 initiator/hairpin system from Molecular Technologies was used for rRNA, while B2 was used for *nosZ* and *ccoN1* mRNA, B3 was used for *narG* mRNA, and B4 was used for *nirS* and *ackA* mRNA. 25 uL of each hairpin were suspended in 20 uL of amplification buffer and added to the cell solutions, and incubated overnight in the dark at 25°C. The cells were then washed three times in 200 uL 5X SSCT, before being resuspended in 20 uL 5X SSCT and spotted onto a microscope slide (catalog number: 3039-002; ThermoFisher) and allowing to dry overnight in the dark at 4°C. Finally, slides were washed with ice-cold ∼0°C ultra-purified water, then with −20°C ethanol, and slides were allowed to air-dry at 25°C in the dark. Each dried cell spot was then covered with 2.5 ml ProLong Gold Antifade Mountant (catalog number: P36930; ThermoFisher), slides were incubated for 48 hours in the dark at 25°C before being stored, and imaged using a Nikon Eclipse Ti2 fluorescence microscope at 40x magnification with 50% laser power and 1 second exposure for the fluorescent channels. Image analysis was performed using FIJI. Phase images were thresholded to produce a mask and detect cells, then the marked cells were thresholded by rRNA signal to eliminate non-bacterial debris from the dataset, then mean intensity was determined for each labeled cell. Quantitative graphs were generated using bokeh.

Hybridization, probe wash, and amplification buffers were acquired from Molecular Technologies (Caltech, Pasadena, CA).

### ABBA Preparation

ABBA samples were prepared as described previously (Jorth, 2020). Two sequential overnight cultures of wildtype *P. aeruginosa* PA14 were grown in the same liquid media as the final ABBAs, shaking aerobically at 37°C, then diluted to OD_500_ .001 in molten LB supplemented with nitrate depending on experimental conditions and 0.5% noble agar. The agar suspension was transferred to chambered cover glass slides and allowed to cool for 10 minutes, before being transferred to a humidified chamber and incubated at 37°C for 12 hours. After 12 hours, blocks were carefully removed from their chambers using a sterilized metal spatula into pre-chilled 1 mL of 4% paraformaldehyde in PBS + ProtectRNA and incubated for 24 hours at 4°C. Blocks were washed three times for 30 minutes each in 1 mL PBS, then incubated for 1 hour in 1 mg/mL lysozyme in 10 mM Tris HCl at 37°C to lyse cell walls, then washed again three times for 30 minutes each in 1 mL PBS and stored in 1 mL PBS. Blocks were checked on a microscope for proper growth patterning (large aggregates at the top and smaller ones in the bottom, for wildtype in LB+40 mM nitrate) prior to continuing. For each condition and each gene target, three biological replicates were performed, in addition to one blank (no cell inoculation) per condition.

### ABBA HCR

For each sample, 500 μL of hybridization buffer was mixed with 1 μL even and odd HCR initiator probes (2 μM stock) for 4 nM ultimate concentration. Each block was incubated in the buffer overnight at 37°C, then washed three times for 2 hours in the 1 ml pre-warmed wash buffer at 37°C. Each sample was then combined with 250 μL of preheated amplification buffer and 5 μL snap-cooled fluorescent hairpins (3 μM stock) for 60 nM final concentration. Samples were incubated overnight in the dark at room temperature, then washed for 6 hours in 1 mL 5X SSCT at room temperature in the dark, then transferred to 500 ul PBS and stored at 4°C in the dark until imaging using confocal fluorescence microscopy. Imaging was performed on a Zeiss LSM 800 in the Caltech Bioimaging Facility. Agar blocks were mounted on a glass-bottomed dish with the original air-agar interface oriented toward the glass bottom. Both the Alexa Fluor 488 rRNA and Alexa Fluor 647 mRNA channels were imaged simultaneously, with a laser power of .2% and a gain of 650 for the 488 channel, and a laser power of 2% and a gain of 750 for the 647 channel. At least four image fields were imaged per sample using a 10x objective with 2x zoom, for a final image field of 319.45 × 319.45 microns. For each image field, a 100 slice z-stack was imaged with an interslice distance of 6.24 microns. Images were framed such that the surface, which appears visible as a plane of rRNA signal due to a lawn of cells growing on the agar surface, was within 4 slices of the top of the image. Quantification of fluorescence signal per aggregate was performed using Imaris v9.5.1. A batch pipeline was created to segment aggregates. The 488 signal intensity of each Z plane was normalized, and then aggregates were segmented based on the thresholded 488 signal, with the threshold chosen manually per imaging date. The normalized signal was only used for the purpose of segmentation, and the mean signal intensity values reported here are based on the raw signal rather than the normalized signal. The mean 488 and 647 mRNA signal intensity, as well as aggregate size, was calculated and exported per aggregate. Depth was defined as distance from the top of the image, and images were cropped for analysis such that the surface was at the top slice using native Imaris functions. Quantitative graphs were generated using seaborn and bokeh. Raw images were displayed using FIJI.

